# TAAR2-9 Knockout Mice Exhibit Reduced Wakefulness and Disrupted REM Sleep

**DOI:** 10.1101/2024.09.09.612114

**Authors:** Sunmee Park, Jasmine Heu, Gavin Scheldrup, Ryan K. Tisdale, Yu Sun, Meghan Haire, Shun-Chieh Ma, Marius C. Hoener, Thomas S. Kilduff

## Abstract

Trace amine-associated receptor 1 (TAAR1) has gained attention for its roles in modulating neural systems, sleep/wake control, and as a therapeutic target for neuropsychiatric disorders. Although TAARs 2-9 were initially identified as non-canonical olfactory receptors, recent studies have identified extra-nasal receptor distribution of multiple TAARs. To evaluate whether TAARs 2-9 have a role in arousal state regulation, we investigated sleep/wake control in male TAAR2-9 knockout (KO) mice. After determination of baseline sleep/wake patterns, the homeostatic response to sleep deprivation and response to TAAR1 agonists were compared between KO and C57BL/6J mice. Although the EEG of TAAR2-9 KO mice had lower power in the delta and theta bands and higher power in the gamma range, sleep/wake states were readily identified. KO mice had more NREM sleep during the dark phase and more REM sleep during the light phase. Sleep/wake was fragmented in KO mice with shorter Wake and REM bouts during the dark phase and more REM bouts during the light phase. KO mice exhibited more REM sleep during a sleep latency test but the homeostatic response to sleep loss did not differ between the strains. A high dose of the TAAR1 agonist RO5256390 increased Wake and reduced NREM sleep in KO mice whereas RO5256390 and the partial TAAR1 agonist RO5263397 suppressed REM sleep. The number of tyrosine hydroxylase-immunoreactive neurons in the ventral tegmental area was significantly elevated in KO mice. These dopaminergic and sleep/wake alterations in TAAR2-9 KO mice highlight the need for further elucidation of the functions of TAAR2-9.

## 1 Introduction

Trace amine associated receptors (TAARs) are a family of G protein-coupled receptors (GPCRs) that are expressed in vertebrates (1, 2). Among these receptors, TAAR1 has been the most extensively studied and shown to modulate neural systems. Expressed in various regions of the central and peripheral nervous systems (3-6), TAAR1 has been implicated in the modulation of dopaminergic, serotonergic, and glutamatergic neurotransmission (6-8) and has emerged as a promising therapeutic target for neuropsychiatric disorders, including schizophrenia and drug addiction (9-12). TAAR1 has also been associated with sleep/wake regulation, as demonstrated in studies utilizing TAAR1 agonists in rats (13, 14), wildtype, knockout (KO) and over-expressing (OE) mice (15, 16), and in non-human primates (17).

Recent structural studies have provided detailed molecular insights into the recognition of endogenous trace amines and drugs of abuse by TAAR1 as well as the selectivity and pharmacology of potential antipsychotic agents targeting TAAR1 such as ulotaront (SEP-363856) and ralmitaront (RO6889450) (18-20). A clinical study showed that, consistent with TAAR1 agonist suppression of REM sleep in animal studies, the TAAR1 agonist ulotaront reduced daytime sleep onset REM periods and nighttime REM duration in patients with narcolepsy-cataplexy (21). However, the magnitude of the effects of ulotaront was not deemed to be clinically sufficient to treat this disorder.

In contrast to TAAR1, the roles of TAAR subtypes TAAR2-9 remain relatively poorly understood but recent studies have demonstrated the presence of TAAR family members in various brain regions. In addition to the original description of the TAAR family which reported TAAR4 expression in the amygdala and hippocampus (1), TAAR2 has been found in the hippocampus, cerebellum, cortex, dorsal raphe, hypothalamus and habenula (22) and TAAR5 expression has been reported in the amygdala, hippocampus, piriform cortex, and thalamic and hypothalamic nuclei (23, 24). With respect to function, TAAR5 has been implicated in adult neurogenesis, dopamine transmission and alterations in anxiety and depressive behaviors (22-25). While the specific functions of these and other TAAR subtypes remain to be fully identified, the expression of these receptors in brain regions associated with monoaminergic neurotransmission, emotion, and cognition suggests their potential involvement in modulating neural activities. Given the intricate interplay among monoaminergic neurotransmitter systems, sleep regulation and psychiatric disorders, investigating the roles of TAARs beyond TAAR1 is of considerable interest.

In this study, we studied TAAR2-9 knockout (KO) mice (26) to elucidate the potential contributions of these receptors to sleep regulation and their interactions with monoaminergic neurotransmitters. Although previously implicated in mating behaviors (26), TAAR2-9 KO mice present a unique opportunity to understand additional functions of these receptors, particularly their involvement in modulating neural activities akin to TAAR1. Given the involvement of TAAR1 in sleep/wake control described above, we evaluated whether TAAR2-9 KO mice exhibit sleep/wake patterns distinct from WT or TAAR1 KO mice. We found that TAAR2-9 KO mice have alterations in non-rapid eye movement (NREM) and rapid eye movement (REM) sleep characterized by increased fragmentation and disrupted sleep architecture. Immunohistochemical analyses revealed a significantly greater number of tyrosine hydroxylase-immunoreactive neurons in the ventral tegmental area (VTA) of KO mice, potentially contributing to the increase in REM sleep duration. This finding suggests a potential link between elevated dopaminergic neurotransmission and the observed differences in sleep patterns. Overall, this study extends the putative involvement of TAARs in sleep regulation and underscores the necessity for further investigations into their functions and interactions with monoaminergic systems.

## 2 Materials and methods

### 2.1 Animals

Cryopreserved embryos of TAAR2-9 KO mice (26) maintained in a pure C57BL/6 background were provided by F. Hoffmann-La Roche Ltd. and resuscitated at the Stanford University Transgenic, Knockout and Tumor Model Center (Palo Alto, CA). After weaning from surrogate mothers, mice were transferred to SRI International where they were maintained on a 12h light/12h dark cycle, with ad libitum access to food and water. A cohort of C57BL6/J male mice (WT; n = 4) bred at SRI International were used as controls; this WT cohort was supplemented by C57BL6/J male mice (n = 9) purchased from JAX mice. Room temperature and humidity were maintained at 21-23°C and 30-50%, respectively, and were continuously monitored. All procedures were approved by the Animal Care and Use Committee at SRI International.

### 2.2 Surgical Procedures

Male TAAR2-9 KO (n = 8) and C57BL6/J (n = 4) mice were subcutaneously implanted with sterilized HD-X02 wireless transmitters (DSI, Inc., St. Paul, MN) for the measurement of electroencephalogram (EEG), electromyogram (EMG), locomotor activity (LMA) and subcutaneous body temperature (T_sc_). Due to signal quality issues, one WT mouse was excluded from the data analysis. As indicated above, the WT cohort was supplemented with another 9 male C57BL6/J mice that were also implanted with HD-X02 transmitters. All mice were approximately 5 months of age at the time of recording.

During surgery, mice were anesthetized with isoflurane, and EMG and EEG leads were routed subcutaneously. EEG leads were inserted through the intracranial burr holes, with one lead over the hippocampal area (+1 mm A/P from lambda, +1 mm M/L) and the ground lead over the cerebellum (−1 mm A/P from lambda, +1 mm M/L). After placement of the EEG leads, dental cement was applied to the skull to affix the wires. EMG leads were sutured to the right nuchal muscle. An analgesia cocktail of meloxicam (5 mg/kg, s.c.) and buprenorphine (0.05 mg/kg, s.c.) was administered for two days post-surgery; meloxicam was administered for an additional day.

### 2.3 EEG and EMG Recording

Mice were acclimated to handling at least seven days before data collection was initiated. Recordings occurred in the animal’s home cage no sooner than three weeks post-surgery, with the wireless transmitters turned on at ZT23 on Day 0 and turned off at ZT24 the following day (Day 1).

Physiological signals collected from transmitters were continuously recorded using Ponemah software (DSI, Inc., St. Paul, MN) and subsequently analyzed.

### 2.4 Experimental Protocols

#### 2.4.1 Baseline EEG and EMG Recording, mMSLT, and 1-h, 3-h and 6-h Sleep Deprivation

After obtaining a 24-h baseline recording for each group, we conducted a murine multiple sleep latency test (mMSLT), consisting of a 20 min sleep deprivation period followed by a 20 min undisturbed sleep opportunity for five consecutive 40 min sessions over a 200 min period (27, 28) beginning at ZT3 (n = 8 KO mice; initially n = 3 WT mice, supplemented by 9 additional WT mice at a later date). One week later, the mice were subjected to sleep deprivation (SD) of 1-, 3-, and 6-h duration ending at ZT6, which was then followed by a 18-h recording. At least 48-h elapsed between each SD experiment.

### 2.5 Pharmacological Studies

One week after the SD experiments, mice were administered *per os* (p.o.) one of the following treatments during the light phase at ZT6, with the experimenter blinded to the dosing condition: Vehicle (10% DMSO in DI water), the TAAR1 partial agonist RO5263397 (1mg/kg), the TAAR1 full agonist RO5256390 (3 and 10mg/kg), or caffeine (10mg/kg) as a positive control. Drug treatments were administered in a balanced order such that all mice received all treatments. To ensure adequate wash-out between each treatment, at least 72-h elapsed between dosings. Mice were recorded for EEG and EMG from ZT23 on the day prior to dosing to ZT12 on the dosing day. The EEG and EMG recordings from ZT6 to ZT12 were manually scored blind to the drug condition and analyzed to determine the effects of the drug treatments on sleep-wake states.

### 2.6 Immunohistochemical (IHC) Studies

Mice (n = 6 TAAR2-9 KO and n = 9 WT mice) were anesthetized with Isoflurane and administered 0.1ml of Somnasol (Pentobarbital sodium, 390 mg/ml). Mice were then perfused with 0.9% NaCl in 0.1 M phosphate-buffered saline (PBS) at pH 7.4 followed by 4% paraformaldehyde in 0.1 M PBS (pH 7.4). After perfusion, the brains were removed and cryoprotected in 30% sucrose solution. The brains were sliced into 40 μm coronal sections at -20°C using a cryotome (Leica CM3050S, Germany). Sections were collected from the ventral tegmental area (VTA) through the dorsal raphe (DR). To identify dopaminergic neurons expressed in these areas, free-floating sections were incubated overnight at 4°C with a polyclonal rabbit anti-tyrosine hydroxylase (TH) antibody (Calbiochem, 657012-100μL) diluted 1:250. Sections were then incubated with a donkey anti-rabbit secondary antibody (Jackson Immunoresearch, 115233) diluted 1:500 for 2 h at room temperature. Slices were then processed with a mixture of diaminobenzidine (DAB), NaCl, and 0.03% H_2_O_2_ (Vectastain DAB kit, Vector Labs, CA). The stained sections were mounted on microscope slides, cover-slipped, and tissue images were obtained using a slide scanner (Olympus VS200, Japan).

### 2.7 Cell Counts

High-resolution images from the slide scanner were analyzed using Fiji ImageJ for cell quantification (29). Images were pre-processed in Fiji ImageJ, adjusting brightness, contrast, and sharpness. Cell identification was performed using a two-step process: first, an initial automated selection based on a set threshold and binary mask creation, followed by manual inspection and confirmation of each identified cell. Cell intensities were evaluated and quantification was performed in the following regions: the ventral tegmental area (VTA), the substantia nigra pars compacta (SNc), the substantia nigra pars reticulata (SNr), and the locus coeruleus (LC). SN sub-areas were delineated according to the Allen Institute’s Mouse Brain Atlas (http://mouse.brain-map.org).

### 2.8 EEG Scoring and Spectral Analysis

Recordings were scored manually in 10-sec epochs as wakefulness, NREM sleep or REM sleep using Neuroscore (DSI Inc., MN) by expert scorers as described previously (15, 16, 28). Wakefulness was characterized by relatively low voltage EEG signals and high EMG. NREM sleep had a higher EEG voltage and dominant power in the delta frequency band but lower EMG. REM sleep had a low EEG voltage as in wakefulness, but with low EMG and dominant power in the theta frequency range. For the baseline and sleep deprivation studies, manually scored data were used as training sets in Somnivore (30) and the full 24-h recordings were autoscored as described previously (31).

EEG spectra for each state were analyzed in 0.122 Hz bins and in standard frequency bands, rounded to the nearest half-Hz value (delta: 0.5–4 Hz, theta: 6–9 Hz, alpha: 9–12 Hz, beta: 12–30 Hz, low gamma: 30–60 Hz and high gamma: 60–100 Hz). Both raw power (Figure 1) and power normalized to the wake state (Figure S1) are presented.

**Figure 1.**
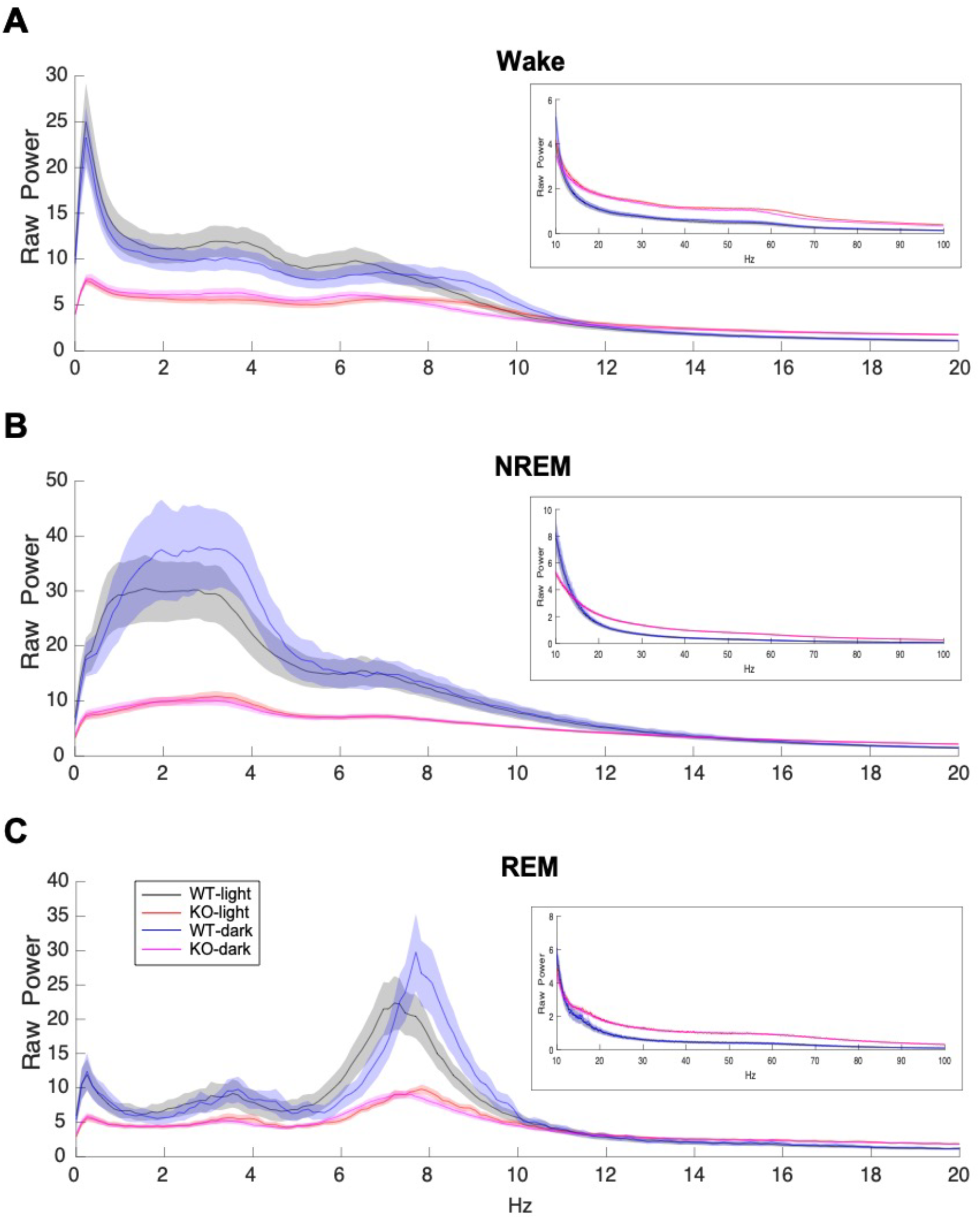
EEG power spectra during Wake, NREM and REM sleep in male C57BL6/J (WT) and TAAR2-9 KO mice during both the light (ZT0-12) and dark (ZT13-24) phases from 0-20 Hz, with insets displaying the 30-100 Hz range. TAAR2-9 KO mice showed reduced EEG power in the low frequency bands in all states during both light and dark phases. However, KOs exhibited higher power in the gamma range (30-100 Hz) compared to WT mice.

### 2.9 Statistical Analysis

Data from the baseline, sleep deprivation and pharmacology studies were analyzed using custom MATLAB R2020b (Mathworks, MA) scripts and statistics functions using Prism 10 (Graphpad, CA).

Images from the IHC studies were analyzed using Fiji (ImageJ for MacOS, opensource package), including the cell auto-counting function. For multiple comparisons, two-way repeated measures ANOVA (RM-ANOVA) was conducted with mouse strain and time (Figures 3-4), drug treatment and time (Figure 6), or the mouse strain and the brain regions (Figure 7) as factors, followed by either Šídák’s or Tukey’s *post hoc* multiple comparison test. For comparisons within the light or dark phases (Figures 3-4) and the baseline vs. sleep deprivation study (Figure 5), the KO and WT groups were compared using an unpaired t-test with Welch’s correction (Welch’s test).

## 3 Results

### 3.1 Baseline Physiological Recordings

#### 3.1.1 EEG Spectral Analysis

The EEG spectra differed considerably between WT and KO mice. KO mice exhibited much lower raw power overall up to 10 Hz but higher power above 20 Hz (Figure 1). These features did not change the overall EEG waveform or the spectral band contribution in KO mice except for the amplitude of the EEG signals. Consequently, Wake, NREM sleep, and REM sleep were readily scored (Figure 2). When analyzed by 2-way ANOVA, normalized power (Figure S1) showed a genotype difference (*p* < 0.0001) during both NREM and REM sleep for all 3-h bins during the light phase. Sidak’s multiple comparisons determined that KO mice had significantly greater normalized power in the β, low and high γ bands during NREM sleep from ZT4 to ZT12 (Figures S1E, S1H and S1K). In contrast, the increase in normalized power during REM sleep progressed with time as only the low γ band was significantly greater from ZT4-6 (Figure S1F), then both the low and high γ bands were significantly greater from ZT7-9 (Figure S1I), and the β, low and high γ bands were significantly greater from ZT10-12 (Figure S1L). Unlike TAAR1 KO mice which had higher normalized power over all bands compared to WT mice (15), no theta band difference was observed in TAAR2-9 KO mice.

**Figure 2.**
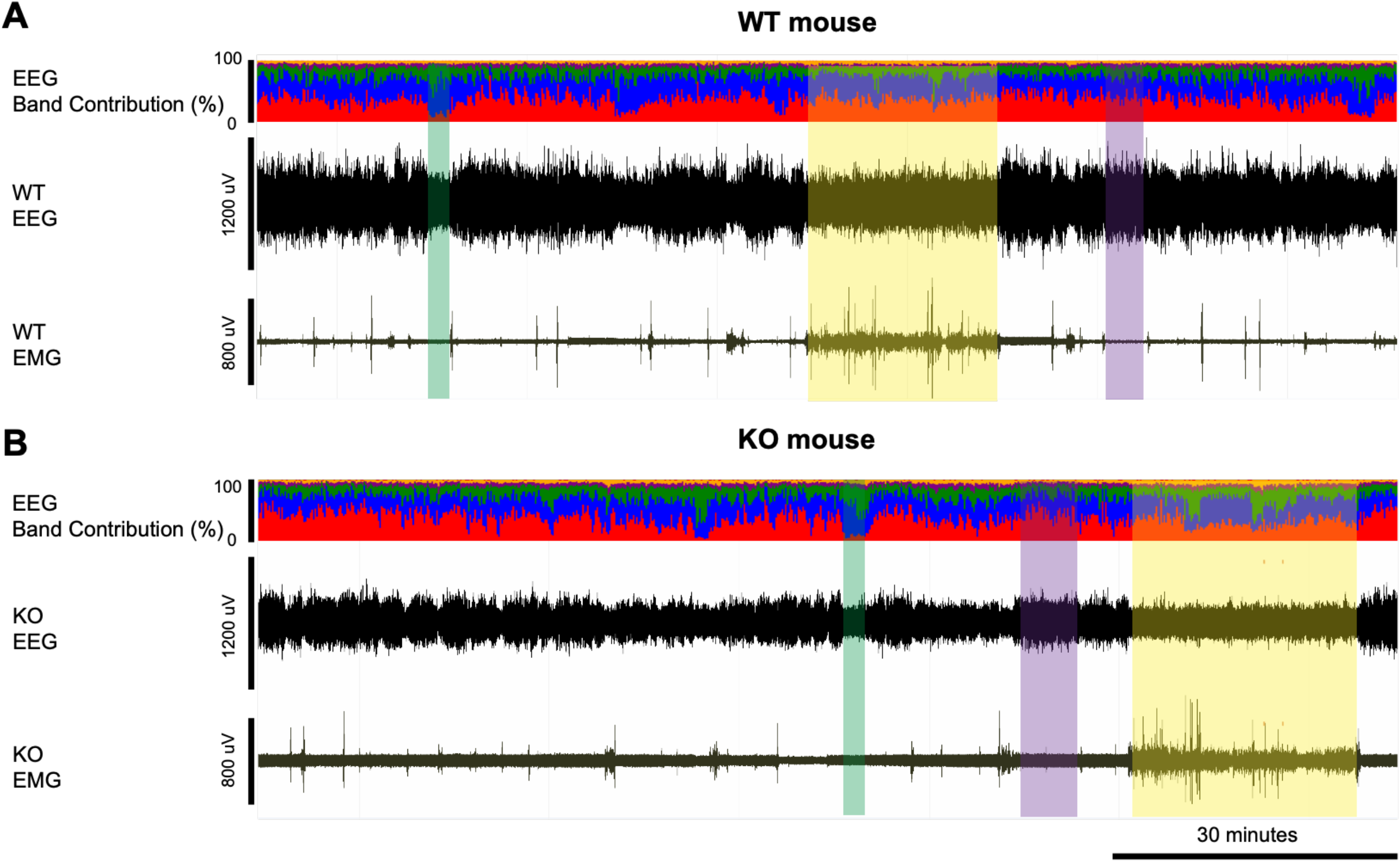
Example EEG and EMG recordings from a (**A**) WT and (**B**) KO mouse. Yellow-shaded boxes indicate periods of wakefulness, purple-shaded boxes indicate NREM sleep, and green-shaded boxes indicate REM sleep. The top trace in panels A and B presents the % contribution of each power band in the EEG signal with colors representing the different frequency bands: delta (0.5-4Hz) in red, theta (4-8Hz) in blue, alpha (8-12Hz) in green, sigma (12-16Hz) in purple, and beta (16-24Hz) in yellow. Wakefulness is characterized by relatively low voltage EEG signals and high EMG while NREM sleep has higher EEG voltage and dominant power in the delta frequency band but lower EMG. REM sleep has similar low EEG voltage as in wakefulness, but with low EMG and dominant power in the theta frequency range. KO mice have relatively low EEG signals compared to the WT mice, as also shown in Figure 1.

#### 3.1.2 Total sleep/wake time and distribution

To assess the sleep/wake patterns of TAAR2-9 KO mice and WT mice, baseline recordings were conducted from ZT0 to ZT24. Figures 3A-C suggested that KO mice exhibited less wakefulness and higher amounts of sleep across the 24-h period. Two-way ANOVA revealed a strong trend for genotype differences in Wake that did not quite reach significance (*p =* 0.0512) and significant genotype (F_(1, 18)_ = 7.719, *p =* 0.0124) and time x genotype (F_(23, 414)_ = 1.857, *p =* 0.0099) differences for REM sleep. Comparison of the total amounts of sleep/wake across the 24-h period by an unpaired t-test with Welch’s correction indicated significantly less wakefulness (*p =* 0.0441) and more REM sleep (*p =* 0.0148) (Figure S2). Analyses of the light and dark phases indicated that KO mice exhibit reduced wakefulness (*p =* 0.0369; Figure 3D) and increased NREM sleep (*p =* 0.0354; Figure 3E) during the dark phase when mice are usually more active. Figures 3A and 3B indicate that these differences were most prominent during the first few hours of the dark phase. During the 12-hour light phase, KO mice also exhibited a significantly higher amount of REM sleep compared to WT mice (*p =* 0.0153; Figure 3F), while no significant differences in the total amounts of Wake or NREM sleep were observed between the two strains at this time of day.

**Figure 3.**
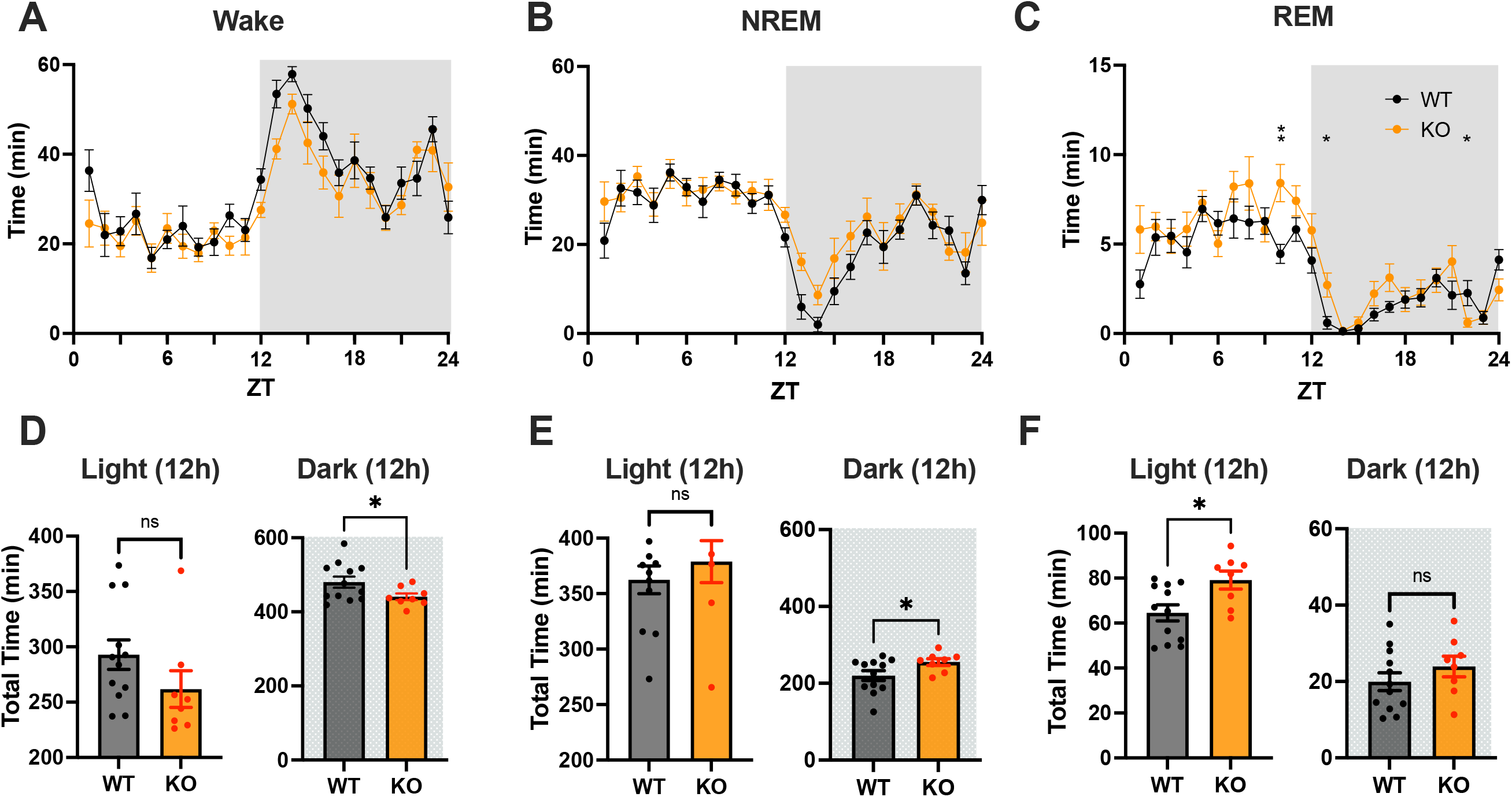
(**A-C**) Distribution of Wake, NREM and REM sleep in male C57BL6/J (WT) and TAAR2-9 KO mice across a 24-h recording. (**D-F**) Cumulative time in each state recorded in each strain during 12-h light (ZT0-12) and 12-h dark (ZT12-24) phases; shaded areas denote the dark phases. WT, Wild type mice; KO, TAAR2-9 KO mice; *, *p* < 0.05; ns, not significant.

#### 3.1.3 Sleep Architecture of KO vs. WT mice

To determine any changes in sleep/wake architecture that underlie the sleep/wake differences between TAAR2-9 KO and WT mice described above, we compared the number of bouts and bout durations between the strains during light and dark phases. The number of bouts per hour and the cumulative number of bouts of Wake and NREM sleep did not differ between WT and KO mice (Figures 4A, B, D and E). However, 2-way ANOVA revealed a genotype difference in the number of REM sleep bouts between the two strains (F_(1, 18)_ = 9.215, *p =* 0.0071; Figure 4C). The cumulative number of REM sleep bouts during the light phase was significantly greater in KO mice (Figure 4F; *p =* 0.0080, Welch’s t-test), although no specific hourly differences were observed.

**Figure 4.**
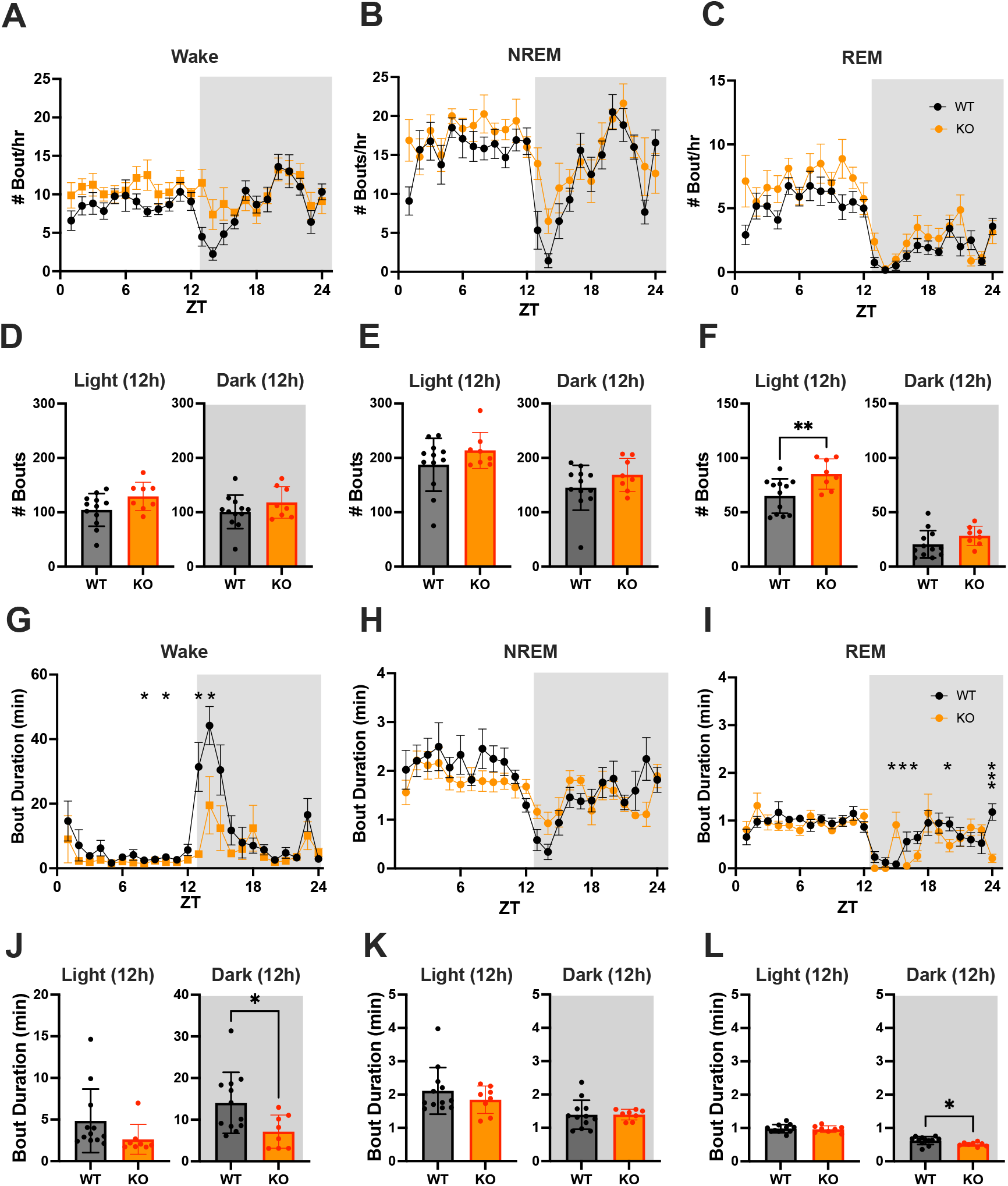
(**A-C**) Hourly number of Wake, NREM and REM sleep bouts in TAAR2-9 KO and WT mice across a 24-h baseline recording. (**D-F**) Mean number of Wake (**D**), NREM sleep (**E**) and REM sleep (**F**) bouts for each strain during the light and dark phases. (**G-I**) Hourly duration of Wake, NREM and REM sleep bouts in TAAR2-9 KO and WT mice across the 24-h recording. (**J-L**) Mean duration of Wake, NREM and REM sleep bouts for each strain during the light (ZT0-12) and dark (ZT12-24) phases. Shaded areas denote the dark phases. Values are mean ± SEM. *, *p* < 0.05; **, *p* < 0.01; ns, not significant.

In contrast, 2-way ANOVA identified both genotype (F_(1, 18)_ = 5.260, *p =* 0.0341) and time x genotype (F_(23, 414)_ = 2.524, *p =* 0.0002) differences for the mean Wake bout duration. Tukey’s multiple comparison *post hoc* test revealed that Wake bouts were significantly shorter in KO mice at ZT8, ZT10, ZT13 and ZT14 (Figure 4G) and Welch’s t-test confirmed that Wake bouts were shorter in the dark phase overall (Figure 4J; *p =* 0.0142), the time of day when mice are normally active.

Two-way ANOVA also identified time x genotype (F_(23, 414)_ = 2.428, *p =* 0.0003) differences for the mean REM bout duration; Tukey’s *post hoc* tests revealed shorter REM bouts in KO mice at ZT15, ZT16, ZT17 and ZT20 but longer REM bouts at ZT15 (Figure 4I). Welch’s t-test confirmed that REM bouts were shorter in KO mice during the dark phase overall (Figure 4L; *p =* 0.0124), suggesting more fragmented sleep in this strain.

These analyses demonstrate that KO mice have disturbed sleep relative to WT mice, particularly fragmented REM sleep with more REM bouts in the light phase and shorter REM bouts in the dark phase. The Multiple Sleep Latency Test (MSLT) further supported these findings, as KO mice spent a significantly higher percentage of time in REM sleep during the recovery sleep (RS) opportunities compared to WT mice (Figure S3E; *p =* 0.0071, Welch’s t-test). KO mice also had difficulty sustaining the long wake bouts that are typical of the early portion of the light phase.

### 3.2 Response to Sleep Deprivation

To investigate the integrity of the sleep homeostatic system in TAAR2-9 KO mice, we evaluated their response to sleep deprivation (SD) durations of 1-h, 3-h, and 6-h compared to WT mice. SDs of these varying duration were initiated to end at the same time of day: the middle of the light phase at ZT6. Each strain’s recording during the SD and subsequent recovery phase was compared to its baseline (BL) day recording (Figure 5).

**Figure 5.**
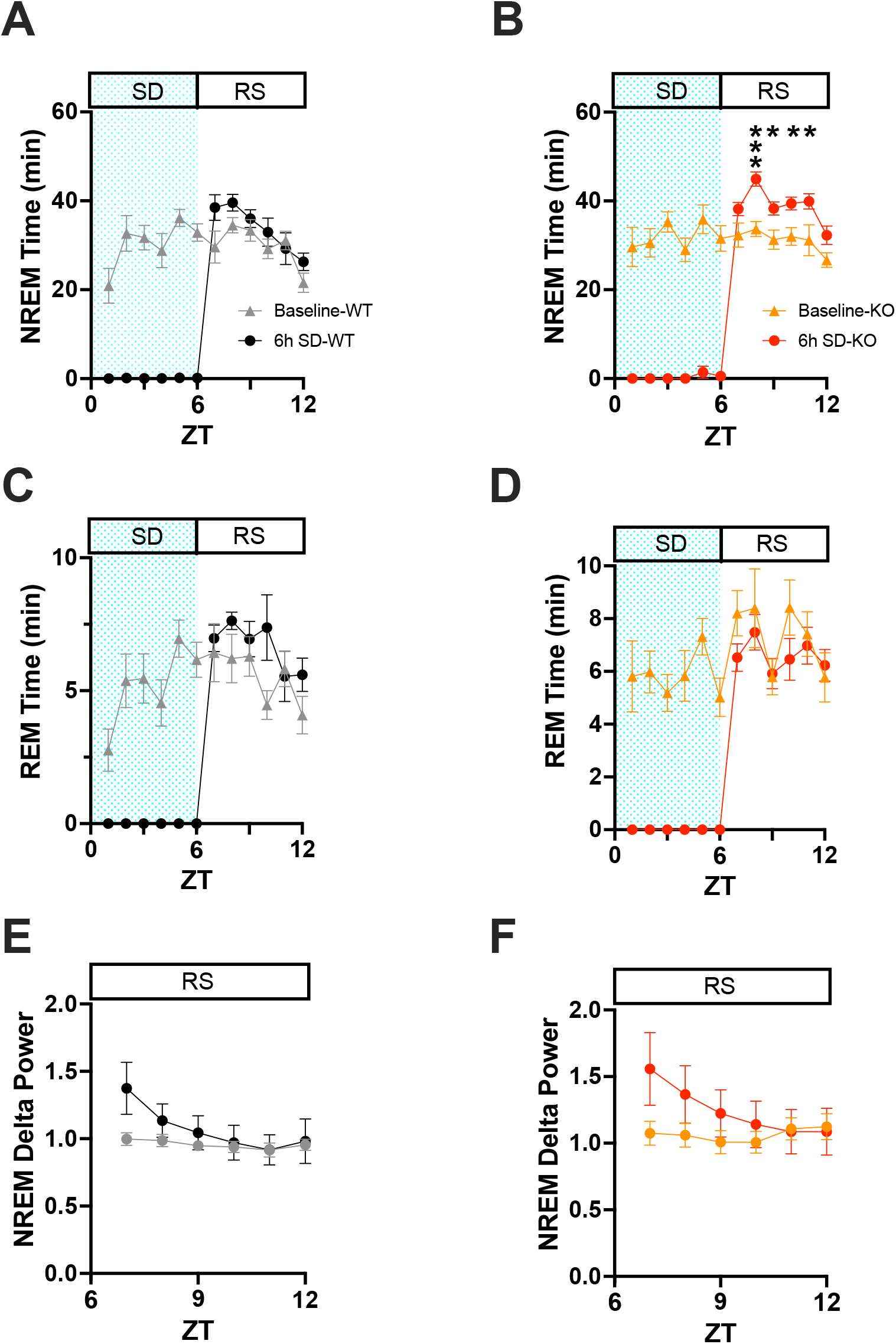
(**A**) Comparison of NREM sleep time from ZT0 to ZT12 in WT mice on a baseline day vs. a day on which 6-h sleep deprivation (SD) occurred from ZT0-6 (shaded area) followed by a 6-h recovery sleep (RS) opportunity from ZT6-12. (**B**) Same as A but for TAAR2-9 KO mice. (**C, D**) Same as A, B but for REM sleep. (**E, F**) EEG delta power in NREM sleep during the RS period after 6-h SD compared to the baseline day in WT (**E**) and TAAR2-9 KO (**F**) mice. Values are mean ± SEM. *, *p* < 0.05.

When the hourly amounts of NREM sleep during the recovery after the 6-h SD were compared to the same ZT6-12 period on the baseline day, 2-way ANOVA found a main effect for condition in both WT (Figure 5A; F_(1, 22)_ = 4.675, *p* = 0.0417) and KO (Figure 5B; F_(1, 13)_ = 20.36, *p* = 0.0006) mice.

However, the NREM increase was significant on an hourly basis only for the KO mice (ZT8-11). Two-way ANOVA also found a condition effect for hourly REM time in WT mice (Figure 5C; F_(1, 22)_ = 4.498, *p* = 0.0455). NREM Delta power during recovery from SD (ZT 7-12) showed time x condition effects in both WT (Figure 5E; F_(5, 110)_ = 8.731, *p* < 0.0001) and KO mice (Figure 5F; F_(5, 70)_= 12.02, *p* < 0.0001), although no specific hour during recovery was significantly different from BL for either mouse strain.

These results from the 6-h SD indicate an intact homeostatic sleep response in KO mice, as exemplified during the first 3 hours of the RS period when NREM delta power increased relative to the baseline day (Figures 5E-F). The results from the 1-h and 3-h SD were similar although not shown here. Both strains showed reduced wakefulness during recovery in the light phase (ZT6-12; Figure S4A), which persisted into the subsequent dark phase (Figure S4B). However, KO mice exhibited more NREM sleep during recovery in both the light (Figure S4C) and dark phases (Figure S4D), suggesting an enhanced homeostatic response to SD. Recovery of REM sleep was delayed until the dark phase in both strains (Figure S4F).

### 3.3 Response to TAAR1 agonists

We tested the TAAR1 partial agonist RO5263397 (1mpk), the full agonist RO5256390 (3mpk and 10mpk) and caffeine (10mpk) to determine whether TAAR2-9 KO mice responded to these compounds. Two-way ANOVA revealed significant effects of drug treatment on Wake (F_(30,30)_ = 2.032, *p* = 0.0264; Figure 6A), NREM (F_(30,30)_ = 1.965, *p* = 0.0346; Figure 6B) and REM sleep (F_(30,30)_ = 2.061, p = 0.0260; Figure 6C). Tukey’s *post hoc* tests determined that caffeine increased Wake (*p* = 0.0115; Figure 6A’) and reduced NREM sleep (*p* = 0.0167; Figure 6B’) during the first 3 hrs post-dosing (ZT6-8) while REM sleep was not significantly affected (Figure 6C’). Compared to Vehicle, the high dose of the full agonist RO5256390 (10mpk) increased Wake for the first three post-dosing hours (Figure 6A, 6A’; *p* = 0.0013) similar to caffeine. During this period, RO5256390 (10mpk) also reduced both NREM (Figure 6B, B’; *p* = 0.0038) and REM sleep (Figure 6C, C’; *p* = 0.0091) The reduction in REM sleep due to RO5256390 (10mpk) persisted into the subsequent 3 hours (ZT8-11; *p* < 0.0001). Interestingly, although RO5263397 (1mpk) had no significant effect on Wake or NREM sleep, it suppressed REM sleep in KO mice during the first three post-dosing hours (Figure 6C, C’, *p* = 0.0324).

**Figure 6.**
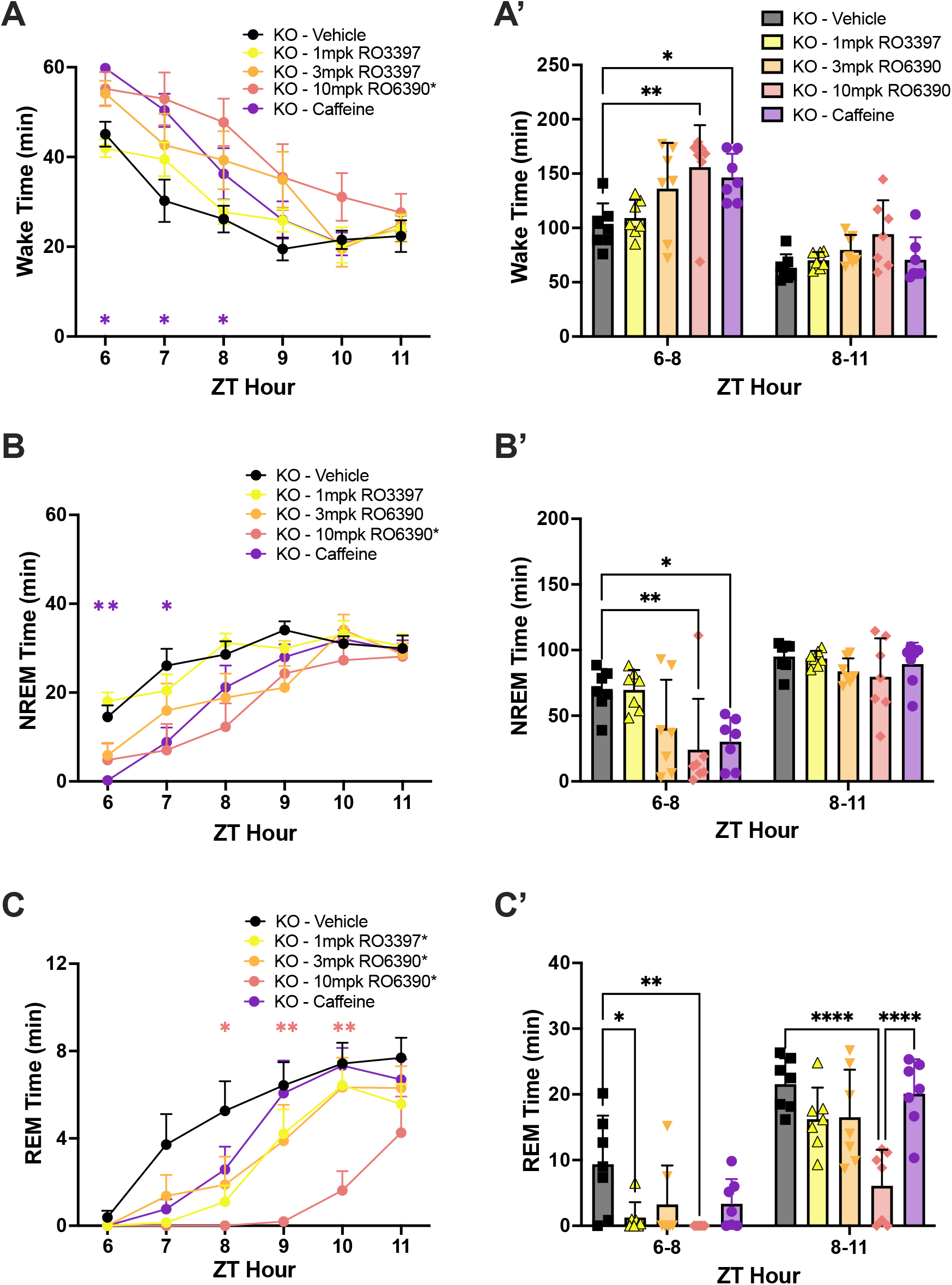
**(A)** Wake, **(B)** NREM, and **(C)** REM time in hourly intervals (left) and summed in 3-hour bins (**A’, B’ and C’**) after dosing with the vehicle, the TAAR1 partial agonist RO3397, the TAAR1 full agonist RO6390 and caffeine. Values are mean ± SEM. Asterisks in the legends indicate a significant condition effect determined by 2-way RM-ANOVA; asterisks within the graphs indicate specific hours when RM-ANOVA indicated a significant time x treatment effect. *, *p* < 0.05; **, *p* < 0.01; ***, *p* < 0.005; ****, *p* < 0.001.

### 3.4 Altered monoaminergic functions in TAAR2-9KO mice

To investigate whether the disrupted sleep patterns observed in TAAR2-9 knockout (KO) mice are associated with alterations in monoaminergic neurotransmitter systems, we conducted immunohistochemical analyses to examine the expression of a key enzyme in dopaminergic neuronal populations. Tyrosine hydroxylase (TH), the rate-limiting enzyme in dopamine synthesis, was used as a marker to label dopaminergic neurons in the ventral tegmental area (VTA), substantia nigra pars compacta (SNc), substantia nigra pars reticulata (SNr), and to identify norepinephrine-producing neurons in the locus coeruleus (LC). Two-way ANOVA revealed significant effects of both Genotype (F_(1,24)_ = 11.76, *p =* 0.0022) and Brain Region (F_(3,24)_ = 32.97, *p* < 0.0001). Tukey’s *post hoc* test determined that TAAR2-9 KO mice exhibited a significantly greater number of TH-positive dopaminergic neurons in the VTA than in wildtype controls (Figure 7; *p =* 0.0019). While a similar trend was observed in the SNc and SNr, these differences did not reach statistical significance. The number of TH-labelled neurons in the LC was similar between the two strains. These findings suggest that the absence of TAAR2-9 receptors may lead to a selective upregulation of the dopaminergic system, particularly in the VTA, a key area involved in dopamine neurotransmission and the regulation of motivation, reward, and motor functions. This selective dopaminergic dysregulation could contribute to the altered sleep architecture and sleep-wake behaviors observed in the TAAR2-9 KO mice.

**Figure 7.**
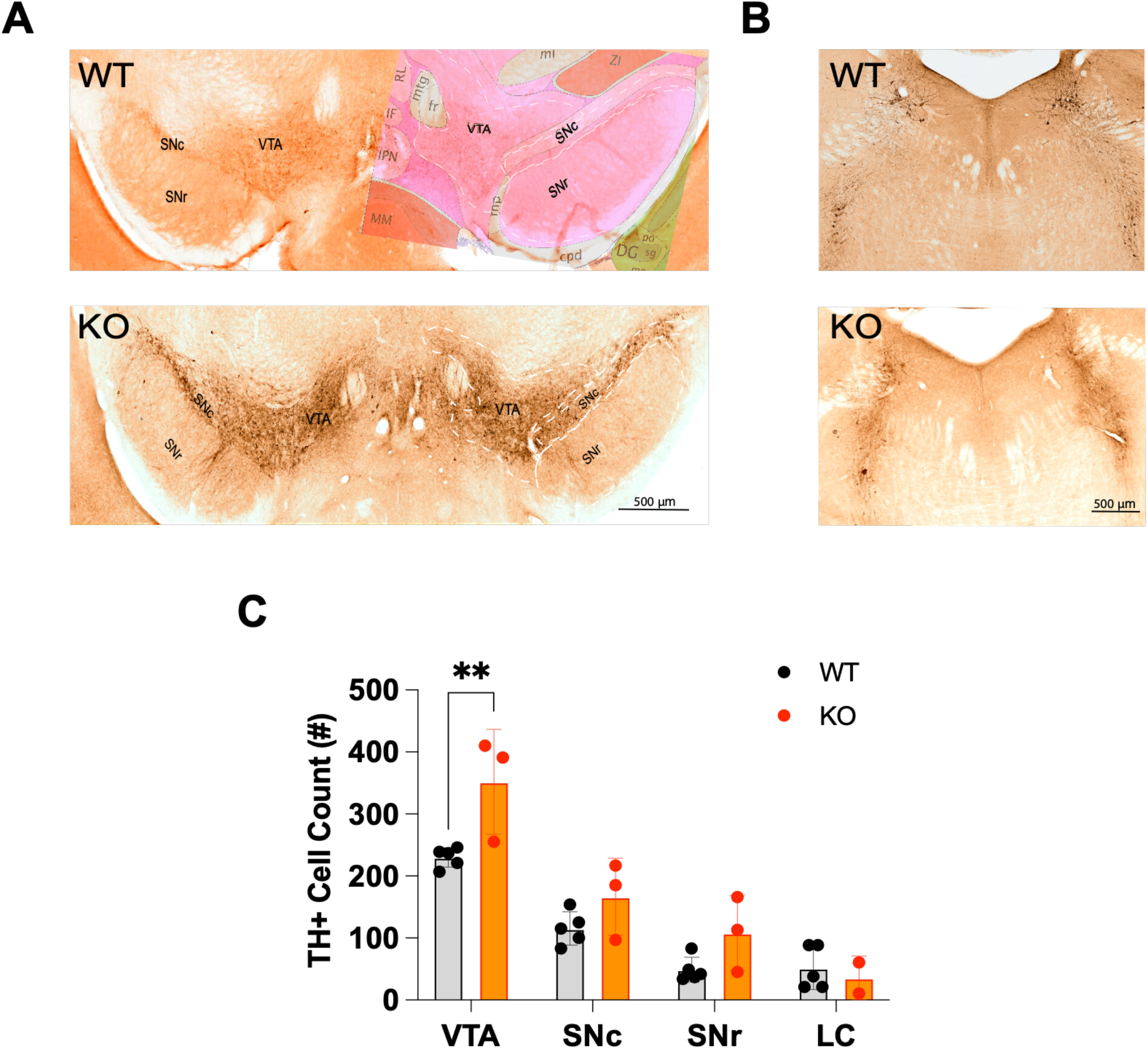
**(A)** Distribution of tyrosine hydroxylase-immunopositive (TH+) neurons in the ventral tegmental area (VTA), the substantia nigra compacta (SNc) and the substantia nigra reticulata (SNr) of TAAR2-9 KO and WT mice. The right side of the WT image in A is overlaid with an image from the corresponding section from the Allen Mouse Brain Atlas. (**B**) Distribution of TH+ neurons in the locus coeruleus (LC) of TAAR2-9 KO and WT mice. (**C**) Number of TH+ neurons in the VTA, SNc, SNr and LC of TAAR2-9KO and WT mice. Values are mean ± SEM. *, *p* < 0.05; **, *p* < 0.01.

## 4 Discussion

The present study provides novel and unexpected insights into the potential roles of TAARs 2-9 in the EEG, sleep/wake regulation and interactions with monoaminergic neurotransmitter systems.

Despite the reduced spectral power in EEG frequencies <10 Hz across all states and increased power in higher EEG frequencies, in conjunction with the EMG, Wake, NREM and REM sleep were readily detected in TAAR2-9 KO mice. Normalization across the entire spectrum revealed that the increased power in the higher frequencies were specific to NREM and REM sleep. TAAR2-9 KO mice had less Wake and more REM sleep across the 24-h period and exhibited distinct alterations in sleep architecture, characterized by reduced Wake and increased NREM sleep during the dark phase, elevated REM sleep during the light phase, and more fragmented sleep patterns overall. TAAR2-9 KO mice were also unable to sustain long bouts of wakefulness early in the dark phase, which is typically the major activity period for mice, and had shorter REM bouts during the dark phase. These observations demonstrate that the absence of TAAR2-9 receptors disrupts the normal sleep-wake cycle and REM/NREM sleep balance and further implicates the trace amine/TAAR system in sleep/wake control as suggested by previous studies (13-17).

Despite the sleep disruptions observed in KO mice, neither sleepiness as measured in the mMSLT nor the homeostatic response to SD as measured in the conventional manner by quantification of EEG Slow Wave Activity (aka NREM Delta Power) differed between TAAR2-9 KO mice and WT controls. However, TAAR2-9 KO mice spent more time in NREM sleep during recovery after SD both during the remaining 6-h of the light phase after SD was terminated and during the subsequent 12-h dark phase. The magnitude of the NREM time increase during the dark phase (and corresponding reduction in Wake time) was significantly greater than in WT mice. These results imply that, in the absence of TAAR2-9 receptors, different compensatory mechanisms may be involved in sleep homeostasis than in WT mice. Further investigations are warranted to elucidate these potential compensatory pathways and their interactions with other sleep-regulatory systems.

We previously reported that the TAAR1 partial agonist RO5263397 increased wakefulness and suppressed REM sleep in rats (13, 14), mice (15) and non-human primates (17). In WT mice, 0.3mpk and 1mpk RO5263397 increased wakefulness and suppressed NREM and REM sleep; those effects were eliminated in TAAR1 KO mice (Table 1), indicating that they are mediated through TAAR1 (15). In contrast, 1.0mpk RO5263397 reduced REM sleep without affecting either Wake or NREM sleep in TAAR2-9 KO mice (Table 1). The suppression of REM sleep by RO5263397 in TAAR2-9 KO mice is consistent with this effect being mediated by TAAR1. However, the absence of an increase in Wake and decrease in NREM sleep in TAAR2-9 KO mice that is observed in WT mice at this dose suggests that TAAR1 may only be partially functional in TAAR2-9 KO mice. One possible source of this discrepancy could be the different genetic background of the TAAR2-9 KO vs. C57BL6/J control mice. Although the TAAR2-9 KO mice were maintained on a C57BL6/J background, the KO and WT control mice used in this study were not littermates.

**Table 1.** Summary of effects of TAAR1 partial and full agonists on sleep/wake parameters in male wildtype, TAAR1 null mutant and TAAR2-9 null mutant mice.

| TAAR1 Compound |  | WT mice |  | TAAR1 KO mice |  | TAAR2-9 KO mice |  |
| --- | --- | --- | --- | --- | --- | --- | --- |
| **Partial agonist** | **RO5263397** | **Sleep/Wake Effect** | **Source:** Black et al. (2017) | **Sleep/Wake Effect** | **Source:** Black et al. (2017) | **Sleep/Wake Effect** | **Source:** Present study |
|  | 0.1 mg/kg | Dec REM only | Fig. 1I | no effect | Fig. 1J, 1K, 1L | not tested | NA |
|  | 0.3 mg/kg | Inc W; dec NREM and REM | Fig. 1G, 1H, 1I | no effect | Fig. 1J, 1K, 1L | not tested | NA |
|  | 1.0 mg/kg | Inc W; dec NREM and REM | Fig. 1G, 1H, 1I | no effect | Fig. 1J, 1K, 1L | Dec REM | Figs. 6C, 6C' |
| **Full agonist** | **RO5256390** | **Sleep/Wake Effect** | **Source:** Black et al. (2017) | **Sleep/Wake Effect** | **Source:** Black et al. (2017) | **Sleep/Wake Effect** | **Source:** Present study |
|  | 1.0 mg/kg | Dec REM only | Fig . 1C | no effect | Fig . 1D, 1E, 1F | not tested | NA |
|  | 3.0 mg/kg | Dec REM only | Fig . 1C | no effect | Fig . 1D, 1E, 1F | no effect | Figs. 6A, 6A'; 6B, 6B'; 6C, 6C' |
|  | 10 mg/kg | Dec REM only | Fig . 1C | no effect | Fig . 1D, 1E, 1F | Inc W; dec NREM and REM | Figs. 6A, 6A'; 6B, 6B'; 6C, 6C' |

In WT mice, the TAAR1 full agonist RO5256390 decreased REM sleep without affecting Wake or NREM sleep at the doses tested (1mpk, 3mpk, and 10mpk); Table 1 summarizes that this effect was eliminated in TAAR1 KO mice (16). By contrast in the present study, the TAAR1 full agonist RO5256390 (10 mpk) enhanced Wake and suppressed both NREM and REM sleep in TAAR2-9 KO mice and the effects on REM sleep were prolonged (Table 1). These results indicate that, in the absence of TAAR2-9, the full agonist RO5256390 produces effects similar to the partial agonist RO5263397 in WT mice. These findings suggest that TAAR2-9 may modulate the effects of these TAAR1 agonists on sleep/wake regulation and suggest interactions between TAAR1 and TAAR2-9, possibly through monoaminergic neurotransmitter systems.

The immunohistochemical analysis indicated an increased number of tyrosine hydroxylase-positive (TH+) dopaminergic neurons in the ventral tegmental area (VTA) of TAAR2-9 KO mice, suggesting a dysregulation of the dopaminergic system in this strain. Given the crucial role of the VTA in dopamine neurotransmission and its involvement in regulating motivated behaviors, reward processing, and motor functions, this dopaminergic alteration may contribute to the observed sleep phenotypes, particularly the increased REM sleep (32).

Overall, these findings contribute to our understanding of the complex interplay between trace amine receptors and monoaminergic neurotransmission in the regulation of sleep-wake cycles and sleep architecture. The observed alterations in sleep patterns and dopaminergic signaling in TAAR2-9 KO mice highlight the potential involvement of these lesser-known TAARs in modulating neural processes underlying sleep regulation.

## 5 Conclusion

The present study provides novel evidence for the involvement of TAAR2-9 receptors in sleep regulation and their interactions with monoaminergic neurotransmitter systems. The altered sleep architecture, disrupted REM/NREM sleep balance, and dopaminergic dysregulation observed in TAAR2-9 KO mice suggests previously unsuspected roles for these receptors in modulating sleep-wake cycles and associated neural pathways. While further investigations are needed to elucidate the precise mechanisms and pathways involved, the findings from this study suggests a new avenue for exploring the therapeutic potential of targeting TAAR2-9 receptors in sleep disorders and neuropsychiatric conditions involving monoaminergic dysregulation. The interactions between TAAR1 agonists and TAAR2-9 receptors, as evidenced by the pharmacological effects on sleep/wake, suggest potential synergistic or compensatory mechanisms that also warrant further exploration. Understanding these interactions may lead to the development of more effective and targeted therapeutic strategies for sleep-related disorders and associated comorbidities.

Overall, this study highlights the importance of expanding our understanding of the lesser-known trace amine receptor subtypes and their contributions to various physiological processes, particularly those involving monoaminergic neurotransmission and sleep regulation.

## Conflict of Interest

The authors declare that the research was conducted in the absence of any commercial or financial relationships that could be construed as a potential conflict of interest.

## Author Contributions

SP and JH conducted all experiments and scored the EEG/EMG recordings. SP and GS performed the immunohistochemistry study. MH, SCM, RT and YS participated in the sleep deprivation study, scoring, and discussion. SP analyzed the data. MCH provided critical reagents for this study. SP and TSK designed the study and wrote the manuscript. All authors reviewed and approved the final manuscript.

## Funding

This work was supported by NIH 1R01NS103529 and 1R01NS136808 to TSK.

## Acknowledgments

The authors acknowledge the assistance of Dr. Hong Zheng, the Stanford University Transgenic, Knockout and Tumor Model Center staff, and the SRI International veterinary care team.

## Notes

### Competing Interest Statement

The authors have declared no competing interest.

