## Supplementary Information for "TAAR2-9 Knockout Mice Exhibit Reduced Wakefulness and Disrupted REM Sleep"

• Supplementary Figure Legends

• Supplementary Figures S1-S4

### Supplementary Figure Legends

**Figure S1.** EEG power spectra during Wake, NREM and REM sleep normalized to Wake for male C57BL6/J (WT) and TAAR2-9 KO mice during the light phase (ZT0-12) recordings. KO mice have higher EEG power in the $\beta$ and both low and high $\gamma$ bands during NREM and REM sleep, particularly during the later 3-hour bins. Values are mean ± SEM. *, $p < 0.05$; **, $p < 0.01$.

**Figure S2.** Total amounts of Wake (A), NREM (B) and REM sleep (C) in male C57BL6/J (WT) and TAAR2-9 KO mice across a 24-h baseline recording. WT, Wild type mice; KO, TAAR2-9 KO mice. Values are mean ± SEM. *, $p < 0.05$; *, ns, not significant.

**Figure S3.** Multiple Sleep Latency Test (MSLT) results from TAAR2-9 KO and WT mice. (A) Percent time in NREM sleep during each 20 min nap opportunity in WT and KO mice. The five 20 min SD periods are indicated by horizontal bars below the abscissa. (B) Percent time in REM sleep during each 20 min nap opportunity. (C) Mean percent NREM time during the 5 nap opportunities in WT and KO mice. (D) Mean NREM sleep latency in the WT and KO mice. (E) Mean percent REM time during the 5 nap opportunities in WT and KO mice. (F) Mean latency to REM sleep between WT and KO mice. In C-F, dots represent the maximum, minimum and 25% and 75% percentile for each bar. Values are mean ± SEM. **, $p < 0.01$.

**Figure S4.** Comparison of the hourly amounts of each state from ZT6-12 on the baseline (BL) day vs. the recording after 6h sleep deprivation (SD) in WT and KO mice. SD data of the light phase is from 6 hours excluding the sleep deprivation (ZT0-6) period and all other periods are from 12 hour of each phase (light: ZT0-12, dark: ZT13-24). (A-B) Wake, (C-D) NREM sleep (E-F) REM sleep. Values are mean ± SEM. *, $p<0.05$; **, $p<0.01$; ***, $p<0.005$; ****, $p<0.0001$.

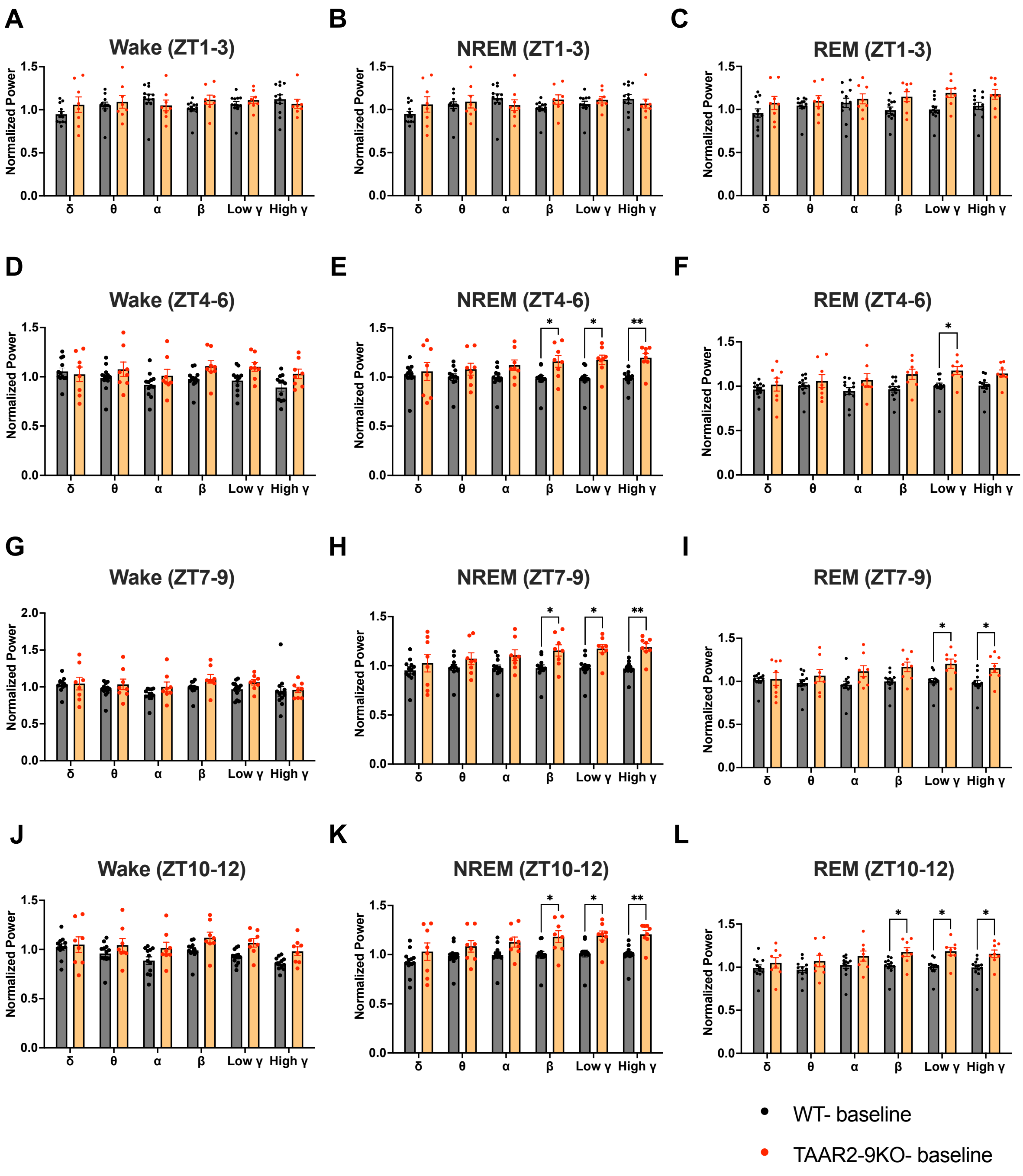
A
Wake (ZT1-3)
B
NREM (ZT1-3)
C
REM (ZT1-3)
D
Wake (ZT4-6)
E
NREM (ZT4-6)
F
REM (ZT4-6)
G
Wake (ZT7-9)
H
NREM (ZT7-9)
I
REM (ZT7-9)
J
Wake (ZT10-12)
K
NREM (ZT10-12)
L
REM (ZT10-12)
Normalized Power
δ θ α β Low γ High γ
WT- baseline
TAAR2-9KO- baseline

**Figure S1.**

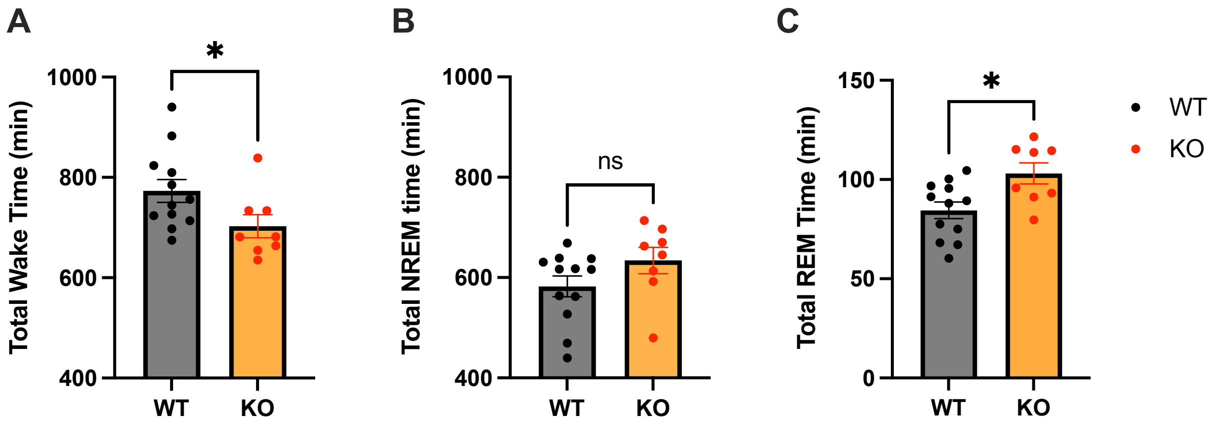
A
Total Wake Time (min)
1000
800
600
400
*
WT
KO
B
Total NREM time (min)
1000
800
600
400
ns
WT
KO
C
Total REM Time (min)
150
100
50
0
*
WT
KO
WT
KO

**Figure S2.**

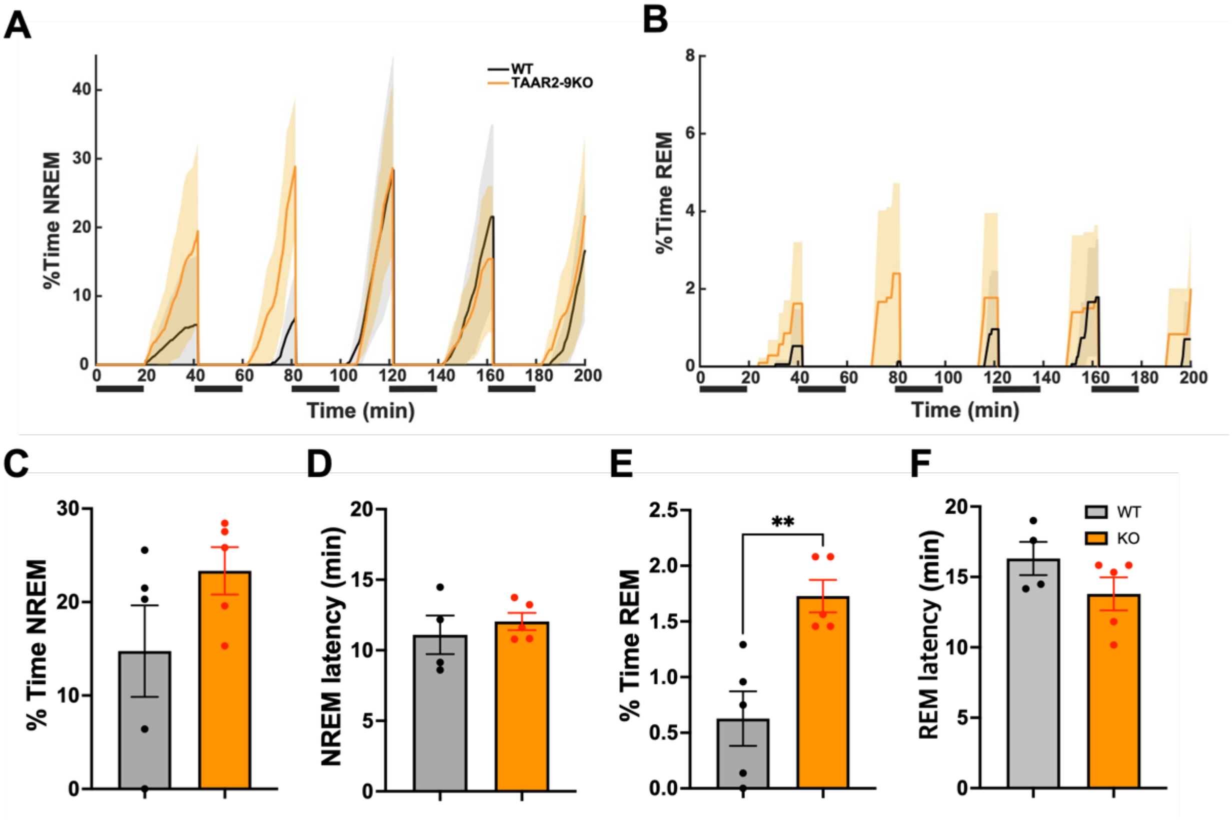
A
WT
TAAR2-9KO
%Time NREM
Time (min)
B
%Time REM
Time (min)
C
% Time NREM
D
NREM latency (min)
E
% Time REM
**
F
REM latency (min)
WT
KO

**Figure S3.**

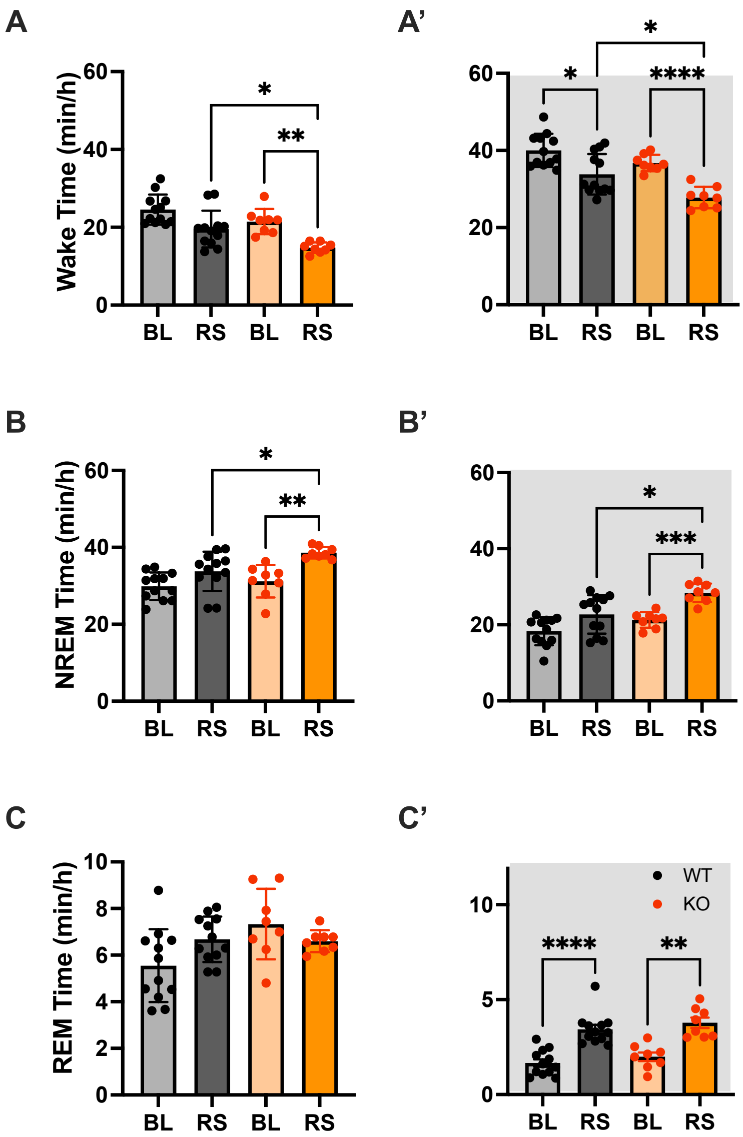
A
A'
Wake Time (min/h)
B
B'
NREM Time (min/h)
C
C'
REM Time (min/h)
BL
RS
BL
RS
WT
KO

Figure S4.
